# Scaling principles of distributed circuits

**DOI:** 10.1101/449447

**Authors:** Shyam Srinivasan, Charles F Stevens

## Abstract

Distributed circuits like the olfactory cortex, hippocampus, and cerebellum contain sub-circuits whose inputs distribute their axons over the entire circuit creating a puzzle of how information is encoded. One method for approaching the puzzle is to view them as scalable systems. In scalable systems the quantitative relationship between circuit components is conserved across brain sizes, and by mapping circuit size to functional abilities - e.g. visual acuity in the visual circuit - scientists have explained information encoding. This approach has not been applied to anti-map circuits as their scalability is unknown. To address this gap in knowledge, we obtained quantitative descriptions of the olfactory bulb and piriform cortex in six mammals using stereology techniques and light microscopy. We found that the olfactory circuit is scalable as it satisfies three requirements of scalable systems. First, quantitative relationships between circuit components are conserved: the number piriform neurons *n* scales with bulb glomeruli *g* as *n* ∼ *g*^3/2^. Second, the olfactory circuit has an invariant property: the average number of synapses between a bulb glomerulus and piriform neuron is one. Third, the olfactory circuit is symmorphic, i.e. olfactory ability improves with circuit size. Other distributed circuits with similar properties might also be scalable.

## Introduction

A potentially powerful way of understanding brain function is by applying the concept of scalability to neural circuits. The approach takes advantage of the fact that developmental constraints often force evolution to conserve a circuit’s architecture across species while varying its size (measured as area, volume, number of neurons, or other quantitative descriptors). By relating the change in size to changes in functional ability, one can relate structure and function. A circuit is considered scalable if the change in size of its components is orderly across species and can be described by simple mathematical relationships such as power laws of the form *Y* ∼ αX^β^, where *β* is the scaling exponent, α is a constant, and *X* and *Y* are size descriptors. Numerous studies have revealed scaling laws for brain circuits to gain insight into circuit function, e.g. [1, 2, 3]. For instance a well known study by Stephan et al. [2] that showed scaling relationships between different regions for insectivores and primates has been cited by more than 700 studies. While a promising first step, these anatomy studies used volumes and areas as proxies for size whereas the brain’s basic computational unit is a neuron. For this reason it is problematic to make inferences about functional abilities and structure.

The number of studies that have used neurons as a measure of size are far fewer, but have yielded profitable results. For example, in the visual system, an increase in the number of visual cortex neurons should correspondingly increase visual ability [4]. These predictions were confirmed in subsequent studies showing that primates have 2.5 times more neurons in V1 than similar sized non-primates, and their visual acuity is about 2.5 times better [5]. But, lessons from the visual circuit might not be universally applicable because brains contain many circuits whose organization is unlike that of the visual cortex.

For the arguments that follow, we need to make a distinction between two types of circuits - topographic and distributed circuits - based on the connectivity of their constituent neurons that we describe below. A ready example of topographic circuits is the retinotopic mapping in the visual circuit from retinal ganglion cells (RGCs) in the eye to the lateral geniculate nucleus (LGN), and from there to primary visual cortex (V1) neurons. When RGCs send information to V1, their cortical targets are predictable (with high probability), e.g. neighbouring RGCs are highly likely to send visual information to neighbouring V1 neurons. This kind of mapping is called topographic. Some circuits, however, are not topographic in the same fashion, but are still predictable, have a mapping between source and target regions. Such topography is observed in the olfactory circuit from the olfactory epithelium (OE) to the olfactory bulb (OB) [6]. Olfactory sensory neuron (OSN) types in the epithelium - uniquely identified by the olfactory receptor gene that they express [7, 8] - respond selectively to odors [9, 10, 11, 12], and each OSN projects to glomeruli in the olfactory bulb that are exclusive to that OSN-type. Thus, even though the OSN targets of neighbours might themselves not be neighbours, they remain predictable. They also display another characteristic of topographic circuits: target regions are highly localized. OSNs of a particular type target one or two glomeruli whose positions are conserved across individuals and occupy a small fraction of the bulb volume [13, 14].

This topographic connectivity contrasts with the connections from the bulb to the piriform cortex (PCx), the biggest brain region subserving olfaction. Here, glomeruli, via mitral and tufted (M/T) cells, contact neurons distributed throughout the piriform cortex without any detectable spatial preference (Fig. 1A,B). Two axons from a glomerulus (or sister M/T cells) are equally likely to target piriform neurons that are neighbours or are far apart. The probability of accurately predicting the targets is no better than if the targets were chosen randomly. In practical terms the targets seem to be randomly distributed, i.e. odor information is distributed throughout the cortex and we term these circuits distributed (or anti-map because of absence of maps) circuits. This distributed organization is reflected in the encoding of odors by a unique combination of neurons distributed over the PCx without any spatial preference [15, ?, 16, 17]. This contrasts with topographic circuits, where information is encoded in the spatial identity of neurons, e.g. bulb glomeruli or V1 neurons that encode a specific OSN-type or point in space. Thus, topographic and distributed circuits differ in the connectivity between source and target neurons, predictability and localization of targets versus unpredictability and distributed nature of targets, and how they encode information. Given these differences, one can imagine that visual circuit scaling relationships might not apply to the piriform cortex.

**Figure 1:**
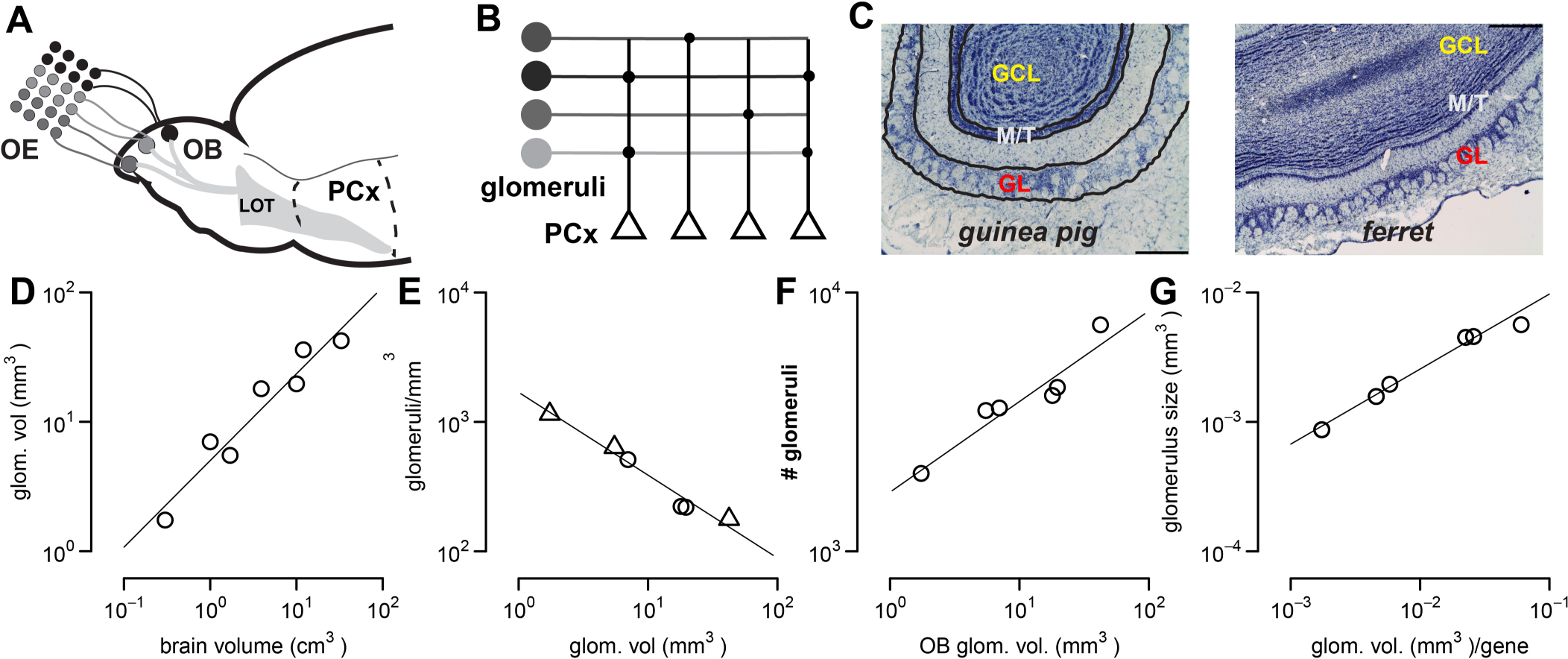
The olfactory bulb and its components increase with larger brains. (A) Schematic of the olfactory circuit in mice. Odor information detected by various types (distinguished by color) of OSNs is conveyed to their cognate glomeruli in the OB, from where it is passed on via the LOT to PCx. (B) A more in-depth schematic of the connections from the glomeruli to PCx neurons. The circles denote synapses which are made without any spatial preference. (C) Representative sections of the olfactory bulb in guinea pigs and ferrets with the glomerular layer (GL), Mitral/tufted (M/T) layer, and Granule Cell layer (GCL) marked. Scale bar: 0.5 mm. (D-G) Scaling of olfactory bulb components. (D) As brain volume (*BV*) increases, so does the volume of the glomerular layer (GV) described by 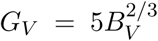, *R*^2^ = 0.93, CI: 0.46 - 0.89. (E) The glomerular density (*g_d_*) reduces with larger glomerular layers or olfactory bulbs according to 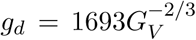. *R*^2^ = 0.97, *CI*: -(0.8 - 0.48). Triangles represent data from the literature and circles represent our measurements. (F) The number of glomeruli increases with bigger bulbs by the relationship 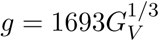, *R*^2^ = 0.9, *CI*: 0.19 - 0.51 (G) As the number of OSNs in the OE denoted by glomerular volume per gene increases, so does the size of an average glomerulus, *g* = 0.0.37*oe*^0.58^ *R*^2^ = 0.98, *CI*: 0.52 - 0.64. Abbreviations: OSN - olfactory sensory neuron, OE - Olfactory Epithelium, LOT - lateral olfactory tract, PCx - piriform cortex, OB -olfactory bulb, CI- 95% confidence interval for the exponent, *R*^2^ - coefficient of determination.

The olfactory circuit is a promising candidate for exploring how distributed systems scale because its function, connection architecture, and response patterns are conserved across phyla [18, 19, 20, 21, 22, 23, 24]. For this reason, olfactory circuits might be scalable as manipulating size is an easy way for evolution to adapt each species to its niche without designing developmental programs, *de novo.* The true test of scalability, however, is if individual components of the olfactory circuit (the olfactory epithelium, bulb, and primary olfactory cortex) change in size such that the relationship between them is conserved across species. To test this hypothesis, we need quantitative estimates of the 3 stages of the olfactory circuit.

In this paper, we provide a quantitative description and test if components of the olfactory circuit -the OE, the olfactory bulb and the primary olfactory cortex - scale across 6 mammalian species. We find that olfactory bulb and piriform cortex sizes increase with more input from the OE. Concomitantly, the number of glomeruli and neurons in Layer 2 (the odor processing layer of PCx), increase while satisfying two relationships. First, the average number of synapses between a glomerulus and a Layer 2 neuron in the piriform cortex is invariant (around 1 synapse) across species. Second, the number of piriform neurons *n* is related to the number of glomeruli *g* as *n* ∼ *g*^3/2^. These two relationships, as described later, ensure that the precision of the odor code is conserved in the olfactory bulb and the piriform cortex, thus scaling the amount of odor information encoded by the circuit and olfactory abilities with size. Together, these findings satisfy the essential characteristics of scalable systems, and provide evidence that the olfactory circuit has a scalable architecture.

## Results

The 3 main components that comprise the olfactory circuit are shown in Figure 1A. We chose 6 mammalian species to examine the scaling of olfactory circuits: mice, rats, guinea pigs, ferrets, cats, and opossums. The species were chosen to account for breadth in terms of brain size (compare mice vs cats, .3 cm^3^ vs 33 cm^3^), coverage of animals from within one clade to observe clade specific trends (mice, rats, guinea pigs), and evolutionary diversity (covering 2 mammalian infraclasses: eutheria or placental mammals versus metatheria or marsupials).

### The number of glomeruli per olfactory sensory neuron type increases with bigger bulbs

In this study, we treat the glomerulus as a computational unit for two reasons. First, glomeruli are sites of integration of OE input shared by all sister mitral cells. Second, sister mitral cell activity across trials and odors, is highly correlated, unlike non-sister mitral cells, and behaves as a coordinated unit [25]. Furthermore, functional studies show that glomeruli associated with a certain behavior are conserved: see pg 24 of [26] for a more indepth discussion or [27, 28] for related discussions in insects and computational analysis. In mice and rats, around a 1000 or 1200 OSN-types contact 2000 or 3600 glomeruli (Table S1), respectively. Thus each OSN-type is represented by two or three glomeruli suggesting a mechanism for increasing information precision with bigger bulbs. More glomeruli per OSN-type means more odor copies and higher precision [29]. As guinea pig, ferret, opossum, and cat olfactory bulbs are bigger than those of rats or mice, we wondered if there might be a similar increase in the number of glomeruli per OSN-type with bigger olfactory bulbs.

To estimate the number of glomeruli in these species, we need to estimate the glomerular density and the volume or surface area of the glomerular layer as the number of glomeruli is the product of these two estimates. We start with volumetric estimates. As brain size increases, the volume of the OB and the glomerular layer increases too (Fig. 1D, S1F). Note that a power-law relationship (*Y* = *αX^β^*) on a log-log plot is a straight line because *Log*(*Y*) = *Log*(*α*) + *βLog*(*X*) wherein the slope is the scaling exponent β. When we compared the increases in bulb volume versus glomerular volumes, we observed that bigger olfactory bulbs (by volume) had bigger glomerular layer volumes, although glomerular layer volume as a proportion of olfactory bulb volume remained the same (Fig. S1D).

Next, we calculated the glomerular density for guinea pigs, ferrets, and opossums. Briefly, we used stereological techniques, and for the sake of consistency we used the same preparations as those used for obtaining mouse, rat, and rabbit glomerular counts (see supplement, section 2 of methods). The glomerular densities are shown in Fig. 1E, and allowed us to estimate the total number of glomeruli in these species as shown in Fig. 1F. The number of glomeruli *g* increased with an increase in olfactory bulb volume as well as glomerular layer volume (*G_V_*), best described by the equation,

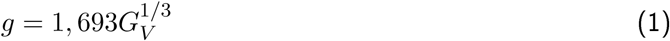

We wondered if the trend that we observed for olfactory bulbs was mimicked by the olfactory epithelium (OE), which is the major input into the bulb [13]. Although we did not measure OE size directly (due to lack of availability), we sought to take advantage of the fact that glomeruli reflect the size of the OE input, and thus serve as a proxy for OE size. Two studies support this assumption. First, a careful study showed that the size of a glomerulus changes with the number of OSNs innervating that glomerulus [30]. A parallel elegant study showed that the number of OSNs of any particular type is environment-dependent because OSN-types that are activated more often, live longer, and occupy a greater fraction of the population of OSNs over time [31]. Thus, as the number of OSNs of any particular type are highly dynamic, glomerular sizes should not be significantly different across different parts of the bulb. This is what researchers have observed [32, 33]. The other consequence is that the average glomerular volume devoted to an OSN type should be a reasonable indicator of the number of OSNs in the OE. We took advantage of these consequence to assess OE and OB size relationships.

With bigger brains, the glomerular layer volume devoted to an OSN-type - a proxy for number of OSNs - increases as shown in Fig. S1E. Thus, bigger brains have more OSNs in the epithelium, correspondingly bigger glomerular layers (Figs. S1F, 1D), and more and bigger glomeruli (Fig. S1G) with two consequences for olfactory coding. The OE input into the bulb (the glomerular volume) and the number of copies of the bulb code (the number of glomeruli per OSN type) increases leading to higher precision of the OE signal and the bulb code.

### Number of Piriform Cortex neurons increases with bigger olfactory circuits

Similar to the OE input into the olfactory bulb, bulbar axons form the biggest input into the piriform cortex. Could the olfactory bulb and piriform cortex (PCx) have a similar scaling relationship? We answered this question by obtaining a quantitative description of the piriform cortex (Fig. 2A, supplement, section 3 of methods) in 6 mammalian species. For ease of explanation, we describe the volumetric and neuronal density estimates, seperately. In all species, the surface area of the piriform cortex increased with brain size (Fig. 2B). We concentrate on the surface area rather than volume as the width of individual layers did not appreciably increase as shown in Fig. S2A.

**Figure 2:**
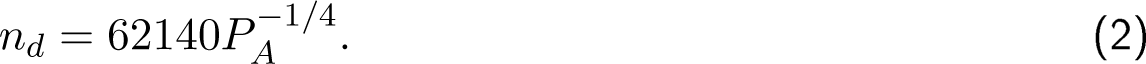
Piriform cortex components increase with bigger brains. (A) Representative sections of the piriform cortex in mice, guinea pigs, and ferrets. The piriform cortex contains 3 layers whose morphological characteristics are clearly distinguishable. Layer 1 is the most superficial layer and abuts the LOT. It is cell-sparse and can be split into Layer 1a, wherein bulbar axons make synapses with L2 and L3 cells, and Layer 1b that contains associative synapses between L2/3 cells. Layer 2 is distinguishable because of the extremely high density of neurons, and is split into Layers 2a and 2b containing semilunar and superficial pyramidal cells. Layer 3 is the deepest layer containing an intermediate number of cells between Layers 1 and 3. (B) The surface area of the piriform cortex (*P_A_*) increases with bigger brains by 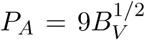. *R*^2^ = 0.95,CI: 0.4 - 0.57. (C) The surface density- number of neurons per mm^2^ of cortex (*n_d_*) - decreases with bigger brains as 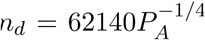. *R*^2^ = 0.84, *CI*: -(0.32 - 0.16). (D) The number of Layer 2 neurons (*n*) increases with bigger brains according to 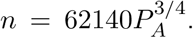. *R*^2^ = 0.98, *CI*: 0.67 - 0.83. (E) The number of L1a synapses increases very gradually with bigger brains as 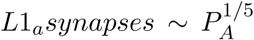. *R*^2^ = 0.54, *CI*: 0.05 - 0.31. LOT- lateral olfactory tract, *L*1_*a*_ -Layer 1a of the piriform cortex, L2/3 - Layer 2/3 of the piriform cortex, PCx - piriform cortex, CI- 95% confidence interval for the exponent, *R^2^* - coefficient of determination.

Next, we obtained surface area density estimates, number of neurons under 1 square mm of cortical surface area, for each layer and the overall piriform cortex for each species (Fig. 2, S2D,E). The surface area density (*n_d_* in *neurons/mm^2^*) decreased with increasing piriform cortex surface area (*P_A_* in *mm*^2^) as shown in Fig. 2C, and given by

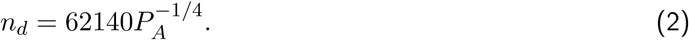

From density and surface area estimates, we calculated the total number of piriform cortex and layer 2 neurons - the main processing layer of the piriform cortex - shown in Figs. 2D and S3C. From here on, when we refer to piriform neurons we mean Layer 2 neurons unless stated otherwise: the results we derive apply equally to the whole neuronal population (Fig. S3C). Overall, as piriform cortices become larger, they acquire more neurons (n) (Fig. 2D), best described by

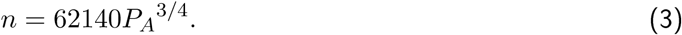

### The piriform cortex scales with olfactory bulb size

Does the olfactory circuit have a scalable architecture? To answer the question, we tested if the relationship between the olfactory epithelium, olfactory bulb, and piriform cortex is the same across species.

Having ascertained that the OE and OB components scale (Figs. ID, S1F-G,S3A), we examined how the piriform cortex changes in relation to the first two stages. We first examined if the surface area of the piriform cortex scales with glomerular volume (Fig. 3A) and found that they are related according to

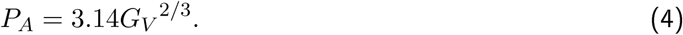

**Figure 3.**
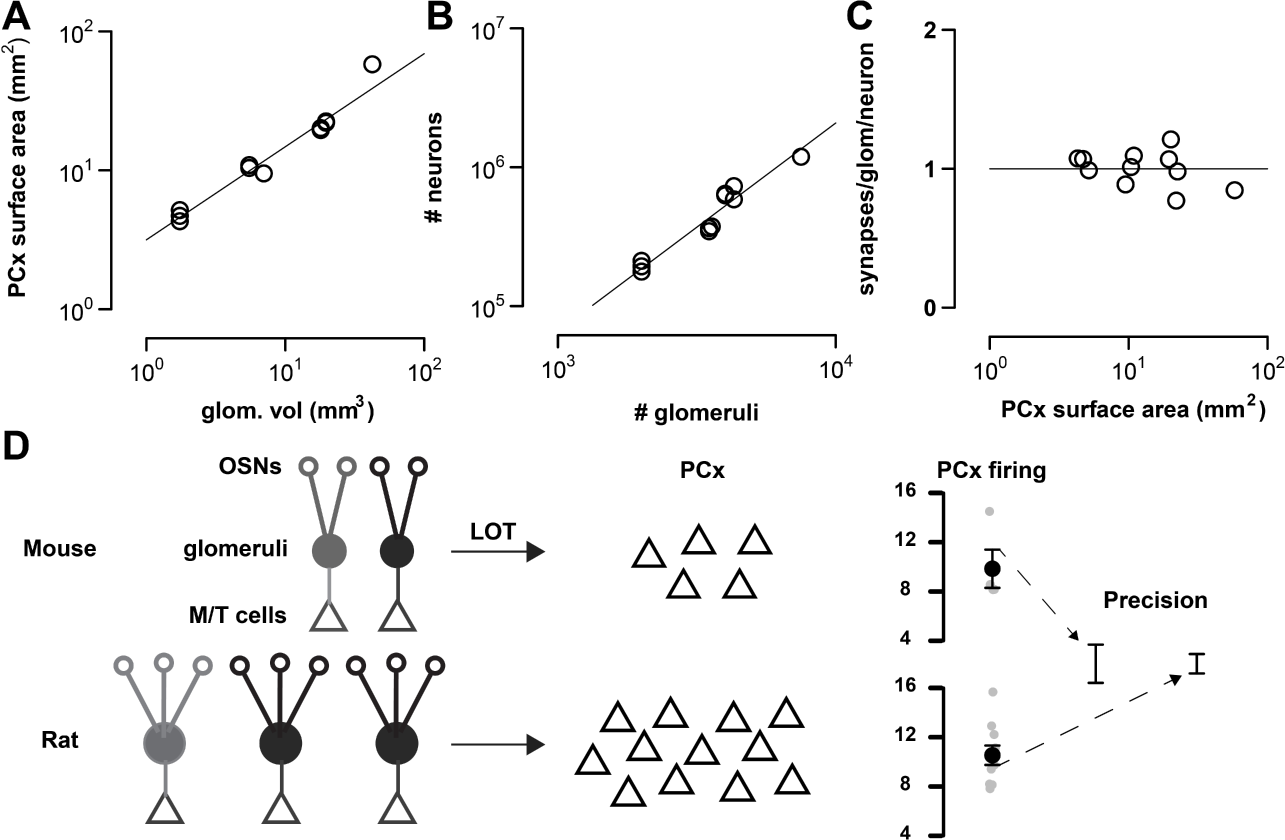
Olfactory circuit components maintain the same relationship as they change in size across species. (A) The surface area of PCx scales with the volume of the glomerular layer (glom. vol) according to 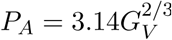. *R*^2^ = 0.96, *CI*: 0.59 - 0.8. (B) The number of neurons in the piriform is related to the number of glomeruli by the relationship, *n* = 2.08g^3/2^, *R*^2^ = 0.93,CI: 1.15 - 1.75 (C) The average number of synapses made by a glomerulus with a Layer 2 neuron is 1 across species (mean = 1.0002, s.e.m = .037). (D) Model of olfactory scaling showing how increasing circuit size improves precision. Compare panels on the top and bottom. As the number of OSNs increases, so does the number of glomeruli as well size of glomeruli. Correspondingly, the number of piriform neurons also increases. The increase in PCx neurons improves the precision of the olfactory code: compare the standard error of the mean (s.e.m) between top and bottom, s.e.m is a measure of the precision of the code. PCx -piriform cortex, CI- 95% confidence interval for the exponent, *R*^2^ - coefficient of determination.

From Figs. 1F (eq.1) and 2D (eq.3), we found that the number of piriform neurons and glomeruli should scale by the relationship, *n* ∼ *g*^3/2^ as we show below in equations (5) - (8).

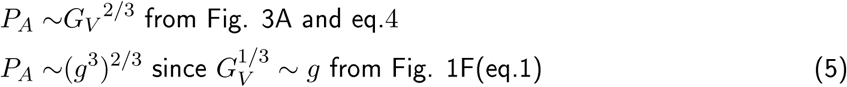

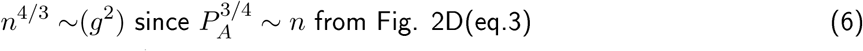

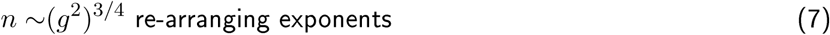

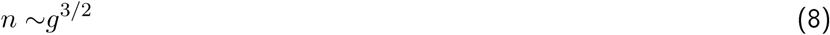

An examination of estimated glomerular and piriform neurons showed that this relationship is indeed true as shown in Fig. 3B (*n* = 2.08*g*^3/2^). Notably, the number of piriform neurons, just like glomeruli also scales with the amount of OE input (Fig. S3B). Overall, olfactory circuit components increase in size with bigger brains, while maintaining the same power law relationship: evidence that the olfactory circuit is scalable.

### Number of synapses between a glomerulus and Layer 2 piriform cortex neuron is conserved

What is the consequence of this scaling relationship on function? To delve into scaling-function relationship, we need a quantitative description of the OB-PCx circuit connectivity. An efficient and parsimonious way to describe connection features of distributed circuits is to calculate the number of synapses that a glomerulus in the olfactory bulb makes with a Layer 2 principal cell in PCx as per Braitenburg and Schuz [34]: here, by a glomerulus’ synapses we mean the synapses made by the M/T cells belonging to that glomerulus.

As there is no spatial preference in the way that OB-PCx synapses are made, the number of synapses is the total number of synapses under a mm^2^ of Layer 1a of PCx divided by number of L2 neurons under 1mm^2^ of PCx and the number of glomeruli in the bulb. In estimating this number, we took advantage of a rich literature that has examined synaptic densities across a wide variety of mammalian species and brain regions and found that it averages to around 1 billion synapses/mm^3^(Fig. S3E, Section 4 and Table S5). We assumed that Layer 1a occupies half the width of Layer 1. The number of Layer 1a synapses under a mm^2^ increases very slightly from the smallest to largest animal (Fig. 2D), and can be expressed as a function of surface area:

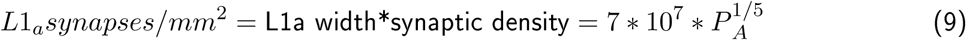

Equations 1 and 3 provide the scaling relationships for glomeruli and neurons. These can be used to calculate the number of synapses (s) between a neuron and - M/T cells of the - glomerulus. It is

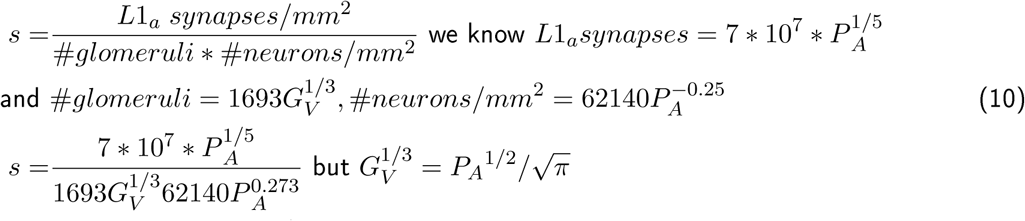

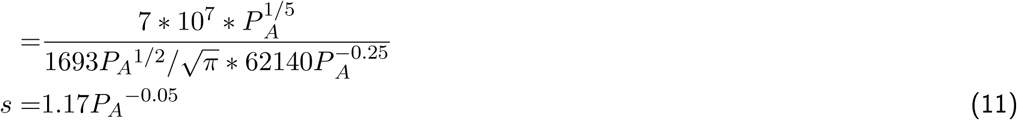

This is close to a constant, and nearly 1, the quantity that emerged from empirical estimates (Fig. 3C). Fig. 3C shows the average number of synapses between a bulb glomerulus and piriform layer 2 neuron, and despite variations in size and ecological niches, we observe a remarkable convergence in the average number of synapses to be around 1 (horizontal line). In the next section, we examine the benefits that such conserved and scaling (eq. 8) relationships might provide.

### Why does the piriform cortex scale with the olfactory bulb?

We provide three reasons why the scaling relationships that we revealed might be advantageous.

**The first reason** stems from the requirement that the precision of the olfactory code should be the same at every stage of the olfactory circuit. Precision is measured as the signal over noise ratio (s/n). As the number of neurons *n* increases, the amount of noise decreases by 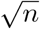, the standard error of the mean. Thus, s/n is proportional to 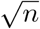. Consider the precision of the odor code at each stage. *N* OSNs of the same type converge onto (*g_OSN_*) glomeruli (*g_OSN_* is the average number of glomeruli per OSN type), which in turn contact *n* neurons in PCx. The precision of the odor code readout from the OE is given by the number of OSNs innervating a single glomerulus: *N/gOSN*- Similarly, all glomeruli of the same type (via their M/T cells) innervate all neurons in PCx, and so the precision of the odor code in the bulb is given by *gOSN,* and the precision of PCx readout is given by *n.* All three quantities must be proportional, which means

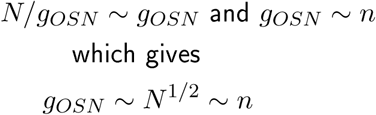

The relationship *g_OSN_* ∼ *N*^1/2^ also follows if we assume optimal allocation of resources in the bulb. If the precision of the OB readout (*g_OSN_*) were to increase, it would reduce the precision of the OE readout *N*/*g_OSN_*- The optimal scenario occurs when (*g_OSN_*) is proportional to *N/g_OSN_* or ∼ *N*^1/2^. On careful consideration, the glomerular volume (*GV*) must be proportional to the number of OSNs innervating it, i.e. *N* ∼ *GV*. In other words, *g_OSN_* ∼ *GV*^1/2^ ∼ *n* which is borne out by our measurements (Fig. S2A, and supplement Sec. 2.2 for a more detailed explanation, including *N* ∼ *GV*). So,

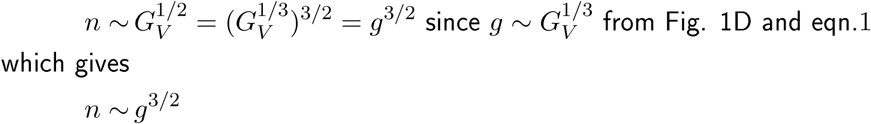

**Second,** we argue that the power-law relationship is required in order for neurons to encode more information with bigger circuits. Each neuron has to make a coding decision based on signals from *g* glomeruli (or from M/T cells belonging to *g* glomeruli), which means the number of inputs (i.e. synapses *s*) to each neuron is *s* ∼ *g*. We also know that the total number of synapses per neuron *s* is, total number of synapses

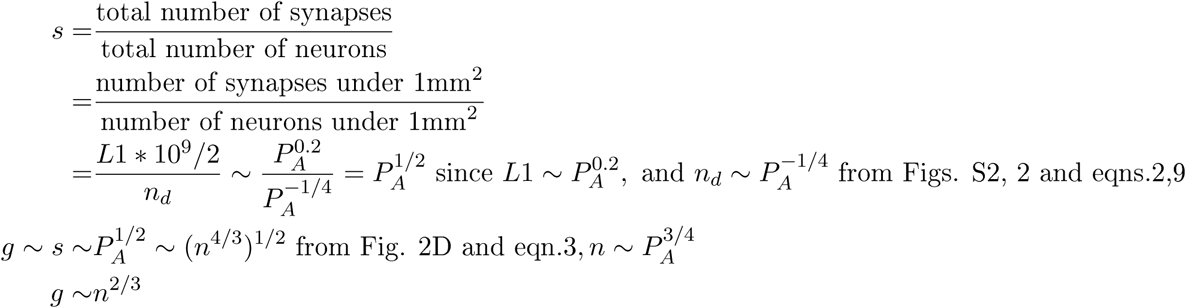

**The third reason** follows from mapping the 2-dimensional (length and glomerular density) bulb system to the 3-dimensional (length, width, and density) PCx system (Section 5 of supplement for explanation). If the output of an 2-dimensional system is to become the input of an 3-dimensional system while maintaining the same precision, the 2-dimensional system should grow faster by 3/2 times, i.e *g*^3/2^ ∼ *n*.

## Discussion

In this study we obtained quantitative estimates of the components within the olfactory bulb and the cortex, and showed that the distributed circuit from the bulb to the piriform cortex, and the topographic sensory circuit from the OE to OB, are scalable. We revealed two principles that olfactory circuits must obey: the average number of synapses between a glomerulus (and its M/T cells) in the bulb and a piriform cortex neuron in Layer 2 is invariant (around one), and the number of neurons *n* in the piriform cortex scales with the number of glomeruli *g* in the bulb asn∼ *g^3^/^2^*-

### Scaling of olfactory abilities

Scaling in biology is not new. It has been observed in myraid contexts ranging from metabolic scaling with body mass in mammals to the scaling of distribution of resources with plant size [35]. Scaling relationships reveal deep design constraints faced by evolution. In every context, scalable systems have been found to possess three properties: the number of (circuit) components increases with bigger (brains) systems such that the relationship between component number and system (brain) size is maintained, a basic system property is invariant across system sizes, and the system is symmorphic, i.e. some quantifiable output of the system increases with size [35]. The olfactory circuit has each of these properties. The number of neurons scales with glomeruli, the number of synapses between a glomerulus and neuron is invariant, and the precision of the olfactory code increases with circuit size, in effect, improving olfactory abilities.

This relationship between improvement in precision and circuit size becomes clear by considering two aspects of olfactory circuit scaling. First, as olfactory bulbs become larger, the number of glomeruli increases (Fig. 2), while the number of olfactory genes does not. This means that the number of glomeruli per OSN-type increases with bigger olfactory circuits, i.e. the number of copies - and precision - of the odor code in the bulb increases. Second, as the precision of the olfactory code is maintained in every part of the circuit, the number of copies and precision of the olfactory code also increases in PCx leading to better detection.

The increase in PCx size will also contribute to better discrimination. Since bulbar axons contact PCx neurons without any spatial preference, their connectivity can be mimicked with a Poisson distribution with a mean of 1. This means that for any glomerulus, there will be a small but significant number of PCx neurons that will receive no synapses, and there will be an even smaller fraction that will receive 5 or more synapses [36]. This fraction could be considered the receptive set for that glomerulus akin to the receptive field in V1, and will differ from the receptive set for other glomeruli. As olfactory circuits acquire more PCx neurons from mice to cats, the receptive set size (or the number of synapses in it) will go up. This difference in receptive set sizes across species will contribute to better discrimination of odors within the piriform cortex.

A natural question that arises is whether behavior data reflects the scaling of ability with circuit size. Behavior studies, however, have provided mixed results [37] because of a few factors. One is the inability to establish a set of odorants that are equally relevant to each of the species being compared. Another is the wide variation in the OSN composition even between individuals of the same species that arise from mechanisms that selectively increase sensitivity to frequently experienced odors [31]. Neverthless, studies showing comparable performance between different species, whose circuit sizes are similar are consistent with the theory [37]. More compelling evidence comes from studies of olfactory circuits during development, aging, disease, and in ecological contexts.

First, in humans, olfactory bulb sizes correlate with odor detection abilities [38]. Second, developmental studies within rodents and humans have shown that as olfactory circuits become bigger, sensitivity and discrimination improve [39, 40, 41]. Third, neurodegeneration caused by age or disease reduces numbers of OSNs and OB volume, and weakens olfactory abilities [42, 43]. Finally, ecologists have shown a positive correlation between bulb and home range size [44, 45]. As the animal ranges further, its olfactory sense needs to improve to sense food over larger distances.

### Topographic circuits vs the distributed olfactory bulb-piriform cortex circuit

Olfactory cortical circuits differ from topographic circuits like the visual circuit in four ways: (1) connection characteristics (seemingly random vs topographic), (2) physiological responses (sparse and randomly distributed ensembles vs topographic), (3) the kind of stimuli they encode (very high dimensional vs few dimensions) and (4) diversity of receptors that sense stimuli. These features are interdependent. A distributed network goes hand in hand with a distributed activation pattern: such patterns cannot arise from topographic inputs. For this reason, many neuroscientists view visual circuit computations as a conduit for understanding other topographic circuits within the neocortex. Some have suggested that computations within a column of the neocortex are similar [46, 47, 48] across species and shared anatomical features such as conserved surface neuron density (number of neurons under a mm^2^) [49, 50] reflect this uniformity.

The quantitative features revealed in this study provide three objective measures by which these neocortical circuits differ from the piriform cortex. First, the number of neurons under a mm^2^ of the piriform cortex is lower (59,000 vs 100,000 in mice) and more variable across species. Second, unlike the neocortex wherein the surface density is conserved across species, PCx surface density decreases with circuit size. The third more subtle feature has to do with the number of inputs innervating the cortices. Neuroanatomists have shown that in the visual cortex, the number of synapses per neuron decreases with bigger brained species [51]. In the piriform cortex, however, the number of synapses per neuron increases with bigger brains. This increase in number of synapses per neuron is accompanied by an equal increase in the number of glomeruli, so that in all species, the number of synaptic connections between (the M/T cells of) a glomerulus and Layer 2 neuron is invariant. These differences add to previous findings in highlighting the divergence between neocortices and piriform cortices suggesting that computational differences arise from architectural differences.

### Evolutionary conservation of distributed circuit features

The same way that scientists view the visual circuit as a prototypical neocortical circuit and a conduit for understanding similar topographic circuits, the OB-PCx circuit could be a conduit for understanding other distributed circuits. Four distributed circuits serve as likely candidates wherein its scaling relationship and connectivity features might be transferable.

The first and the best described one is the fly olfactory circuit. In the fly, high dimensional odor features are captured by OSNs communicated to the mushroom body (PCx analogue), through glomeruli in the antennal lobe (olfactory bulb analogue). Here too the connection matrix from glomeruli to mushroom body neurons is seemingly random [50] and odors are represented by a sparse and random neuronal ensemble [52]. Although differing in circuit details, such homology between the small fly and mammalian circuits suggests that distributed connectivity might be a common motif of distributed and sparse coding.

The second circuit with distributed features is the mammalian olfactory circuit during development [41]. From a young age, even before their olfactory circuit has reached its full size, animals rely on their sense of smell. They need a fully functioning olfactory circuit, which also means that these developing systems should conform to the scaling relationships that we uncovered. Such analogy between developing and adult systems is not new. Previous studies have shown that the ratio of inhibitory to excitatory cells in the neocortex is maintained from a very early developmental stage highlighting the need for inhibitory-excitatory balance in brain functioning [53].

Finally, sparse and distributed ensemble coding is also a feature of the cerebellum and the hippocampus [54]. Studies suggest that the connectivity in these circuits mirrors the distributed connectivity from the olfactory bulb to cortex. If future studies confirm such connectivity, scaling principles from the olfactory circuit might apply to these regions too.

## Methods

The methods that are a part of the supplement include the experimental procedures for histology and stereology, statistical tests for stereology, animal details, and specifics of the counts for glomeruli and PCx neurons.

## Acknowledgments

We would like to thank Saket Navlakha, Joseph Zak, and Terry Sejnowski for valuable feedback on the manuscript. We are grateful to Leah Krubitzer, James Dooley and Andrew Halley for Opossum brains, E.J. Chichilnisky, Edward Callaway, Clare Hulse, Kristina Nielson, and Fumitaka Okasada for Guinea pig, Rat, and Ferret brains. We are also grateful to the Kavli Institute for Brain and Mind at UCSD and NSF-1444273 for supporting this work.

